# Antimalarial and *Plasmodium falciparum* serpentine receptor 12 targeting effect of a purinergic receptor antagonist FDA approved drug

**DOI:** 10.1101/2020.12.16.423179

**Authors:** Sonal Gupta, Nishant Joshi, Monika Saini, Shailja Singh

## Abstract

Intraerythrocytic stages of *Plasmodium falciparum* responsible for all clinical manifestations of malaria are regulated by array of signaling cascades and effector molecules that represents attractive targets for antimalarial therapy. G-protein coupled receptors (GPCRs) are druggable targets in treatment of various pathological conditions however there is limited understanding about the role of GPCRs in malaria pathogenesis. In *Plasmodium,* serpentine receptors (PfSR1, PfSR10, PfSR12 and PfSR25) with GPCR like membrane topology have been reported with the finite knowledge about their potential as antimalarial targets. Herein we evaluated the druggability of PfSR12 using FDA approved P2Y type of purinergic signaling antagonist, Prasugrel. Blocking of purinergic receptor signaling resulted in inhibition of growth and development of *P. falciparum*. Progression studies indicated the inhibitory effect of purinergic signalling inhibitor begins in late erythrocyte stages predominantly in the schizonts. Furthermore, purinoreceptor inhibitor blocked *in vivo* growth of malaria parasite in mouse experimental model. The localization of PfSR1, PfSR10, PfSR12 and PfSR25 was analyzed by immunofluorescence assays. The putative purinergic receptor PfSR12 is found to be expressed in late intraerythrocytic stages. Prasugrel, a generic purinoreceptor inhibitor and agonist ATP showed specific binding to recombinant PfSR12 confirming it as purinergic receptor. This study indicates the presence of P2Y type of purinergic signaling in growth and development of malaria parasite and suggests PfSR12 as putative purinergic receptor.

## Introduction

Malaria remains a ruinous infectious disease caused by Plasmodium sps. *P. falciparum* is one of the most deadly among all five species of Plasmodium known to infect humans. Lack of effective vaccine and increasing resistance to existing antimalarials give way towards the development of novel strategies to combat the disease. Signal transduction pathways play central role for targeting many pathological conditions. Deciphering new signal transduction pathways in *P. falciparum* would be a fundamental step to combat malaria. Intraerythrocytic stages of parasite life cycle are associated with the clinical manifestations of the disease during which Plasmodium merozoites invade erythrocytes through a complex process involving multiple specific interactions between host receptors and parasite ligands ^1^. The role of second messengers such as Ca^2+^ and the downstream signaling cascades has been extensively studied in regulation of invasion phenotype ^2^. However limited knowledge is available about membrane receptors that regulate the signaling machinery in malaria parasite.

In eukaryotic systems, extracellular signaling via nucleotides like ATP, ADP occurs by binding of these nucleotides to specific membrane receptors known as purinergic receptors and controls several biological processes ^3^. Purinergic receptors are broadly of two types: P1 and P2 receptors. P1 receptors belongs to GPCR family of proteins and are activated by adenosine while P2 receptors, further categorized into P2X (ion gated channels) and P2Y (G protein coupled transmembrane receptors) are activated by different nucleotides like ATP, ADP etc ^4^. Purinergic signaling via these receptors leads to upregulation of second messenger like cytosolic calcium and cAMP in the cells ^3^. Under normal physiological conditions, erythrocytes release ATP which functions as signaling molecule that maintains vascular tone ^5^. Some recent reports indicated the significance of extracellular ATP in growth and invasion of erythrocytes by Plasmodium via purinergic signaling ^6,7^. It has been shown previously that purinergic receptor blocker Suramin inhibits secondary processing of merozoite surface protein1 (MSP1) and merozoite invasion ^8^.Together these studies emphasize upon importance of purinergic signaling in malaria pathogenesis. The molecular identification of purinergic receptor in Plasmodium will pave way towards development of novel antimalarials.

*P. falciparum* contains four serpentine receptors (SR) proteins with the seven transmembrane structures like GPCRs namely SR1, SR10, SR12 and SR25 ^9^. Some studies reported the role of subset of these serpentine proteins like SR1, SR10 in salivary gland sporozoites as putative receptors that participate in intracellular signalling cascades regulating parasite’s motility and invasion in host ^10^. Few studies suggested the role of these receptors in development of parasite in host erythrocytes ^11,12^. Bioinformatics analysis of *P. falciparum* serpentine receptor 12 i.e. PfSR12 (PF3D7_0422800) have shown consensus P-loop motif in the protein which could serve as binding pocket for ATP. Based on structural prediction and its strong binding to ATP, PfSR12 is predicted to be a putative purinergic receptor ^13^. Collectively, these studies underline the fundamental importance of serpentine receptors in parasite biology making them promising drug targets. Here, we show the plausible role of P2Y type of purinergic signaling in malaria pathogenesis, where PfSR12 could be identified as putative purinergic receptor in parasite and novel drug target for development of antimalarials.

## Material and Methods

### Reagents

RPMI 1640 medium, Albumax I and gentamicin sulphate (Invitrogen, USA), sodium bicarbonate, Sorbitol, Methanol, Giemsa, SYBR Green-I, Glutaraldehyde, Prasugrel (no. SML0331; Sigma Aldrich-Merck,USA), Suramin (no. S2671; Sigma Aldrich-Merck, USA), ATP (Sigma Aldrich-Merck, USA), ADP (Sigma Aldrich-Merck, USA), R138727 (procured from Biocell Corporation, India), RIPA buffer (no. R0278; Sigma Aldrich-Merck, USA), Hypoxanthine, Alexa-Fluor 488 conjugated goat anti-mouse IgG antibody, Alexa-Fluor 594 labelled goat anti-rabbit IgG, ProLong Gold antifade reagent with DAPI (4’, 6-diamidino-2-phenylindole) (Life technologies-Invitrogen, USA), ATP Affinity kit (no. AK-102, Jena Biosciences).

### In vitro parasite culture

*Plasmodium falciparum* 3D7 was cultured as per the standard protocol ^14^. Briefly, the culture was grown in complete RPMI 1640 medium (RPMI 1640 medium with 2 mM l-glutamine, 25 mM HEPES) supplemented with 2 g/l NaHCO_3_, 50 mg/l hypoxanthine and 0.5% Albumax I, pH7.4 using O^+^ human RBC. Parasite culture was maintained in mixed gas environment (90% N_2_, 5% CO_2_ and 5% O_2_). The parasites were cultured under these conditions in 75 sq cm culture flask. Parasites were synchronized in consecutive two cycles by sorbitol treatment at ring stage. Culture parasitemia levels were monitored by Giemsa-stained smears under 100 X by light microscopy.

### FACs Analysis

The growth inhibition assay (GIA) was performed as described previously using modulators of purinergic signalling. Briefly, synchronized trophozoites at 0.5 % parasitemia and 2 % hematocrit were transferred in flat bottom 96 well tissue culture plate and incubated with different concentrations of purinergic receptor inhibitors like Prasugrel (human P2Y_12_ receptor antagonist) and Suramin (P2 purinoreceptor antagonist) for one growth cycle of parasite with the harvest at trophozoite stage (i.e. 48 hours). The parasite-infected erythrocytes were stained with 10 μg/ml ethidium bromide dye in PBS for 30 minutes and the parasitemia was measured by fluorescence-activated cell sorting (FACS) using BD Fortessa flow cytometer. The parasitemia levels were estimated by determining the number of FL-2 positive cells in the sample. Excitation wavelength of 488 nm was used and about 50,000 cells per sample were scored for fluorescence. For all flow cytometry experiments initial gating was done with unstained RBCs to minimize the background fluorescence. The growth inhibition was calculated with respect to untreated control as following:

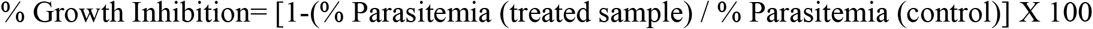

### Fluorescence assay using SYBR Green

To determine the functional relevance of P2Y receptor antagonist Prasugrel in *P. falciparum* growth in human erythrocytes, we performed the growth inhibition assays by SYBR Green method as described earlier ^15^. Synchronized ring stage parasites(~ 12hours post invasion) at 0.5 % parasitemia with a final 2 % hematocrit were incubated with different concentrations (1-50 μM) of Prasugrel: FDA approved drug and P2Y receptor inhibitor) for one growth cycle till trophozoite stage i.e. 60 hours post treatment. Prasugrel pharmacologically active metabolite, R138727 was also tested at concentrations 10 and 20 μM. The assay was terminated by freezing at −80 °C and erythrocytes were lysed by adding lysis buffer containing SYBR green (0.2 μL of SYBR green I/mL of lysis bufer) to each well and incubated for 2 hours in the dark at 37 °C. The fluorescence intensity was determined at 485 nm excitation and 530 nm emission using a Varioskan Flash multi-well plate reader (Thermo Scientifc). The percent growth inhibition was calculated with respect to untreated control and readings were normalized with respect to uninfected RBCs. The concentration of Prasugrel inhibiting parasite growth by 50 % (IC_50_) was estimated by plotting values of percent inhibition against log concentration of compound at different concentrations, using Graphpad PRISM software.

### Light microscopy

Thin blood smears of parasitized erythrocytes; Control and treated cultures were made at different time points of asexual blood stages. Slides were fixed in methanol, air dried and stained with Giemsa stain and examined by light microscopy. The images were captured using CatCam camera and processed by Catymage software (Catalyst Biotech).

### Progression assay

The effect of Prasugrel on parasite development was determined in asexual blood stages. For progression assay, 10-12 h ring-stage of parasite was diluted to 1 % parasitemia and 2 % hematocrit in complete RPMI medium. The culture was treated with Prasugrel at 10 μM and incubated at 37° C for 48 hours. The parasitemia at each stage was calculated by counting around 5000 total erythrocytes in Giemsa stained smears at 1000X magnification using light microscope.

### In vivo studies

Female Swiss mice (n=8), 4-6 weeks age weighing about 25-30 g were used to determine antimalarial effect of FDA approved drug Prasugrel on malaria infection in mice. All protocols were carried out in accordance as per standard guidelines of international animal care and welfare. 1X 10^6^ *P*. *berghei* ANKA infected erythrocytes were injected in the mice by intraperitoneal route. The mice were randomly distributed in two groups (n=4) i.e. vehicle control and Test group. Test compound i.e Prasugrel (stock in DMSO) in 1X PBS was administered in mice by intraperitoneal route at dose of 5 mg/kg body weight. The effect of drug on parasitemia was determined by preparing thin smears of blood obtained from tail vein and stained with Giemsa stain. The parasitemia was estimated by counting 3000-5000 red blood cells in light microscope.

### Expression and Purification of recombinant proteins

The nucleotide sequences encoding amino acids of N-terminal region of PfSR1, PfSR10, PfSR12 and PfSR25 (Plasmo DB accession no. PF3D7_1131100, PF3D7_1215900, PF3D7_0422800 and PF3D7_0713400 respectively) were cloned in pGEX4T1 vector. The recombinant proteins rPfSR1_47_, rPfSR10_44_, rSR12_40_ and rSR25_35_ were expressed in *E. coli* BL21 Rossetta cells with GST tag for affinity purification. Briefly, the bacterial cells were transformed with the expression plasmids and cultured in Luria broth at 37° C. Protein expression was induced at a culture optical density of 0.6-0.7 at 600 nm (OD_600_) with 0.8 mM isopropyl--D-thiogalactopyranoside (IPTG) for overnight. The cell biomass was resuspended in lysis buffer (140 mM NaCl, 2.7 mM KCl, 10 mM Na_2_HPO_4_, 1.8 mM KH_2_PO_4_, pH 7.4) containing 1 mM PMSF, 1 mM Dithiothreitol (DTT) and 100 μg/ml Lysozyme. The supernatant was loaded on to pre-equilibrated Glutathione Sepharose 4B beads (GE Healthcare) and recombinant proteins were eluted by different concentrations of glutathione (5-40 mM). The recombinant proteins were characterized by SDS-PAGE and Coomassie staining.

### Western Blot Analysis

Protein fractions were separated on 12 % SDS-PAGE gel under reducing conditions, transferred to nitrocellulose membrane, blocked with 5 % powdered skimmed milk, and probed with primary antibody (anti PfSRs polyclonal sera raised in mice) at a dilution of 1:500 for one hour at room temperature. The membrane was incubated with HRP conjugated anti-GST secondary antibody (Sigma –Aldrich, USA) at 1:5000 dilution for one hour. After PBS-T washing, the blots were visualized using enhanced chemiluminescent system (ECL).

### Immunofluorescence assay

Thin smears of parasite cultures at different blood stages were made on glass slides, air dried and fixed in pre-chilled methanol. After blocking with 3 % BSA in PBS, the slides were incubated with specific PfSRs mouse sera at 1:50 dilution along with MSP1 (rabbit sera at 1:50 dilution) for co-localization studies. The slides were washed with PBS with 0.05 % Tween-20 and incubated with secondary antibodies like Alexa fluor 488 conjugated anti-mouse IgG goat sera at a dilution of 1:200 or Alexa fluor 594 conjugated anti-rabbit IgG at a dilution of 1:500. Finally, the slides were mounted with ProLong Gold antifade reagent with DAPI (4’, 6-diamidino-2-phenylindole). The images were visualized using Olympus confocal microscope and processed by Imaris software (Bitplane Scientific).

### ATP Affinity Assay

To assess the binding of ATP and Purinergic receptor antagonist Prasugrel with recombinant protein rSR12_40_, ATP binding assays were performed according to the manufacturer’s protocol (Jena Biosciences). Here we used 6AH-ATP-Agarose (N6 - (6-Amino) hexyl-ATP-Agarose) beads in which ATP is immobilized to the beads via adenine base and phosphate group is free. Blank agarose beads were taken as negative control. Briefly, the protein sample was incubated with pre-equilibrated ATP–agarose beads for 2-3 hours at 4° C with slight agitation. For competition assays, the beads were pre-incubated with ATP, ADP or antagonist at desired concentrations. The bead pellets were given stringent washings to remove any unbound proteins with 1X wash buffer. The protein bound to the ATP agarose beads was eluted by ice cold elution buffer containing 0.2 mM sodium orthovanadate and 1 mM DTT and all the fractions were analyzed by SDS-PAGE.

### Molecular docking

PfSR12 protein sequence was taken from PlasmoDB database (PF3D7_0422800). To generate the Ab-initio model of PfSR12 the I-TASSER server was used and validated as described earlier ^13^. Generated model was further subjected to structural refinement by using ModRefiner ^16^. The refined 3D structural model of PfSR12, thus generated, was subsequently used for docking studies. Chemical structures of molecules were synthesized through the ChemSketch ^17^. Ligands structure were optimized using ChemBio3D ultra 12.0 ^18^. Autodock version 1.5.7RC1 and Cygwin terminal was utilized to execute the docking commands ^19^. The binding site for the ligand was chosen around the protein. PLIP and Pymol 2.3.2 softwares were used for further analysis and visualization of docking results ^20,21^.

## Results

### Pharmacological modulators of Purinergic signalling inhibit growth of P. falciparum

Purinergic signaling in malaria parasite has been shown to be involved in invasion of host red blood cells ^6–8^. Prasugrel, a known P2Y_12_ receptor antagonist and FDA approved drug, is a potent inhibitor of platelet activation and aggregation and thereby used in the treatment of thrombotic disease such as coronary heart disease ^22^. We examined the antimalarial activity of Prasugrel by growth inhibition assay in *P. falciparum 3D7.* As shown in Figure 1A, Prasugrel inhibited parasite growth in a dose dependent manner with IC_50_ of 9.6 μM. Pharmacokinetics studies with Prasugrel in humans have shown that this prodrug is activated to pharmacologically active metabolite R-138727 that inhibits platelet aggregation *in vivo* ^23^. Assuming that the antimalarial effect is due to transformation of Prasugrel to active metabolite *in vitro,* we tested inhibitory activity of active metabolite at Prasugrel IC_50_ value i.e.10 μM. We also selected IC_70_ concentration of Prasugrel corresponding to 20 μM, which was showing saturating inhibitory activity. Our results showed R-138727 reduced parasite growth by only 27 % as compared to ~ 75 % inhibition in Prasugrel treated parasite at same concentration 20 μM (Figure 1B). This suggested Prasugrel’s active metabolite has good antimalarial activity but it is required in higher concentration to show similar effect as prodrug. The reduced availability of active metabolite could be attributed to its degradation and instability in experimental set up. Further we confirmed IC_50_ of Prasugrel by flow cytometry in comparative study using known purinergic antagonist Suramin. The results indicated that as compared to Suramin which showed ~ 75 % inhibition at 100 μM, Prasugrel drastically reduced parasite growth with > 90 % inhibition at 20 μM concentration (Figure 1C). At 10 μM concentration of Prasugrel, parasite growth was inhibited by 40 %, which is similar to IC_50_ of Prasugrel by SYBR Green assay.

**Figure 1.**
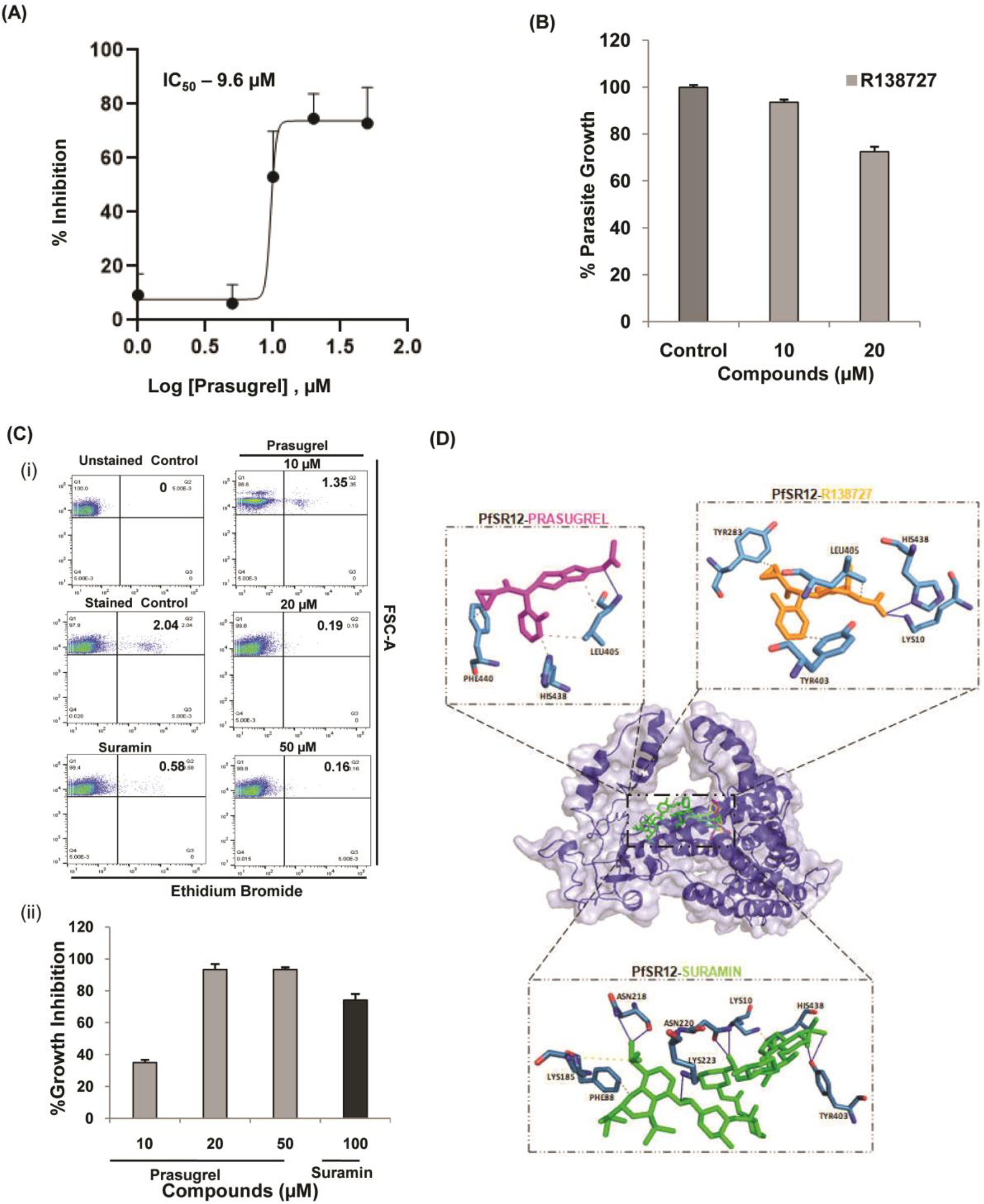
Effect of FDA approved drug and potent purinoreceptor antagonist Prasugrel on *P. falciparum* growth. (A) To calculate half maximal inhibitory concentration (IC_50_) of Prasugrel, *P. falciparum* 3D7 culture at ring stage was treated with different concentrations (1-50 μM) for one growth cycle. IC_50_ value was determined by plotting values of percent inhibition against log concentration of Prasugrel. IC_50_ value is calculated to be 9.6 μM. (B) The growth inhibition by Prasugrel was compared with R-138727 (Prasugrel active metabolite). Bar graph represents significantly high growth inhibition by Prasugrel at 20 μM as compared to R-138727, which shows ~ only 27 % inhibition at same concentration. The experiments were performed in triplicates, n=3 and ± SD value was calculated for each data point.(C) (i) The growth inhibition by Prasugrel was compared with Suramin (non selective purinergic antagonist). Bar graph representing growth inhibitory effect of pharmacological modulators of purinergic signaling on *P. falciparum* (ii) Representative dot plots at each concentrations of inhibitors were shown. Unstained infected erythrocytes were taken as control for the experiment. (D) 3D surface representation of the docking analyses of PfSR12 with purinoreceptor antagonists Prasugrel, Suramin and R138727. Box shows the interacting residues of the complex.

*In silico* studies predicted that all purinergic inhibitors including Prasugrel, R138727 and Suramin interacts with PfSR12, which is previously shown to be a putative purinergic receptor (Figure 1D) ^13^. Together our results indicate that prodrug Prasugrel and Prasugrel like molecules could be good antimalarials. Our findings indicate the plausible role of P2Y type of purinergic signaling in the growth and development of *P. falciparum.*

### P2Y purinergic receptor antagonist Prasugrel arrests parasite growth at schizonts stage

During the intraerythrocytic development, Plasmodium divides mitotically and metamorphoses into different stages namely rings, trophozoite and schizonts in host erythrocytes. Rupture of late schizonts releases 16-20 merozoites in bloodstream which then invade fresh erythrocytes and initiate new cycle of growth and multiplication. To test the effect of Prasugrel on malaria parasite *in vitro* maturation and propagation in erythrocytes, we performed progression assay with untreated and Prasugrel-treated synchronized *P. falciparum* cultures. As shown in the light micrographs of Figure 2, untreated cultures displayed healthy parasites with the normal 48 hours life cycle, with progression from ring to trophozoite and schizont stages of development to reinvasion within the expected time. In contrast, 10 μM Prasugrel-treated cultures showed interference in normal parasite development, with arrested schizonts observed prominently at the 48 hours post treatment. Moreover, many infected erythrocytes showed dead parasites with condensed DNA. Overall parasitemia in untreated control has reached to ~ 3.6 % as compared to ~ 1.9 % parasitemia in Prasugrel treated culture after one cycle. These data suggest that exposure to P2Y inhibitor Prasugrel inhibits progression of malaria parasite in their normal asexual stages.

**Figure 2.**
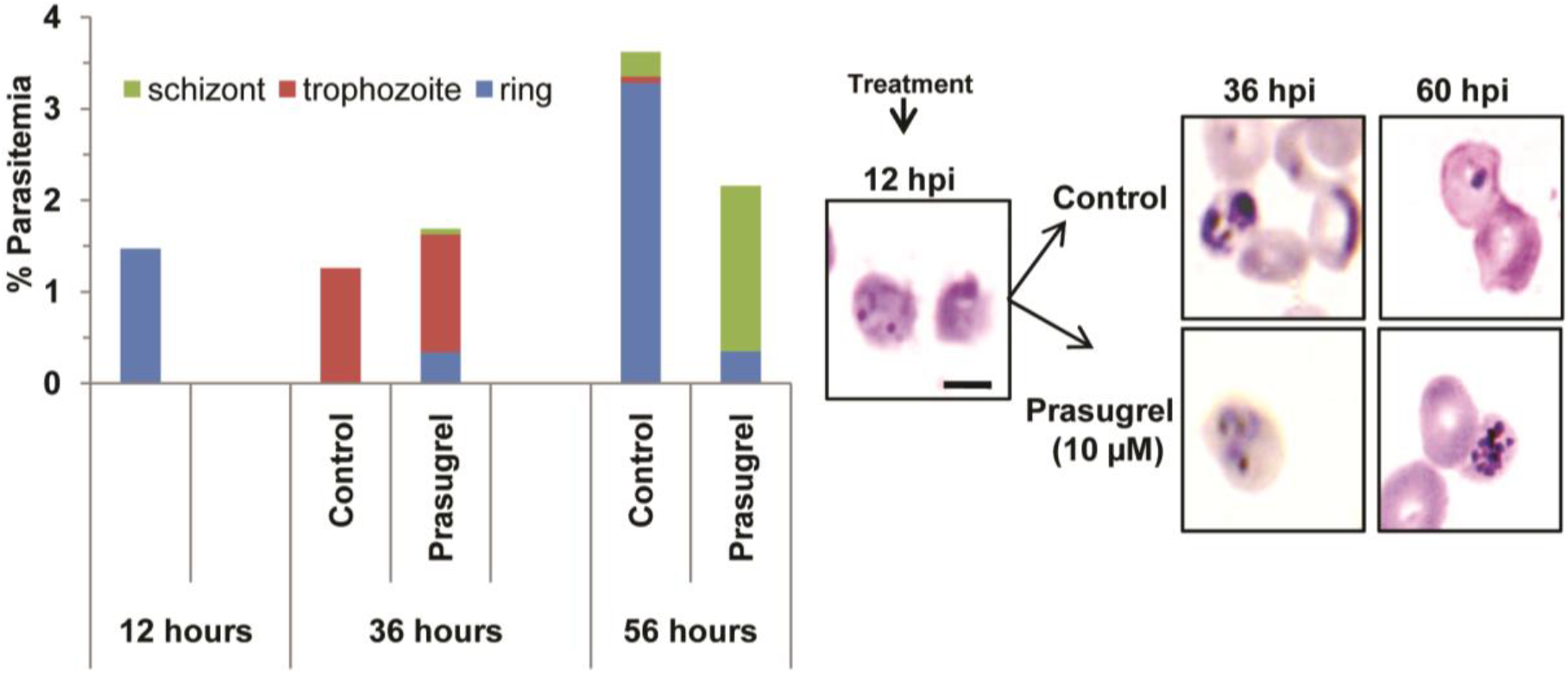
Effect of Prasugrel on the progression of *P. falciparum* 3D7 through intraerythrocytic stages. Synchronous *P. falciparum* cultures were treated at ring stage (12 hpi: hours post invasion) with Prasugrel at IC_50_ concentration (10 μM) and morphology of parasites was assessed by Giemsa stained smears at different time intervals (24 h and 48 h post treatment) i.e. 36 hpi and 60 hpi as shown (Panel on right). Parasitemia quantification was shown by the bar graph for each of the time point for Prasugrel treated and untreated culture. Note that expected ring stage parasites are observed in untreated control after one developmental cycle whereas Prasugrel treated culture stalled at schizonts stage. Scale=5μm.

### Prasugrel blocks in vivo parasite growth

Furthermore, we investigated the effect of P2Y type purinergic signaling on the growth of malaria parasite *in vivo* using rodent malaria model *P*. *berghei.* At Day 7 post infection in mice, the parasitemia showed significant increase to about 30 % in control as compared to only 3 % parasitemia in Prasugrel treated mice confirming the antimalarial effect of the drug (Figure 3). Our *in vivo* results corroborates with growth inhibitory potential of Prasugrel as shown by *in vitro* studies.

**Figure 3.**
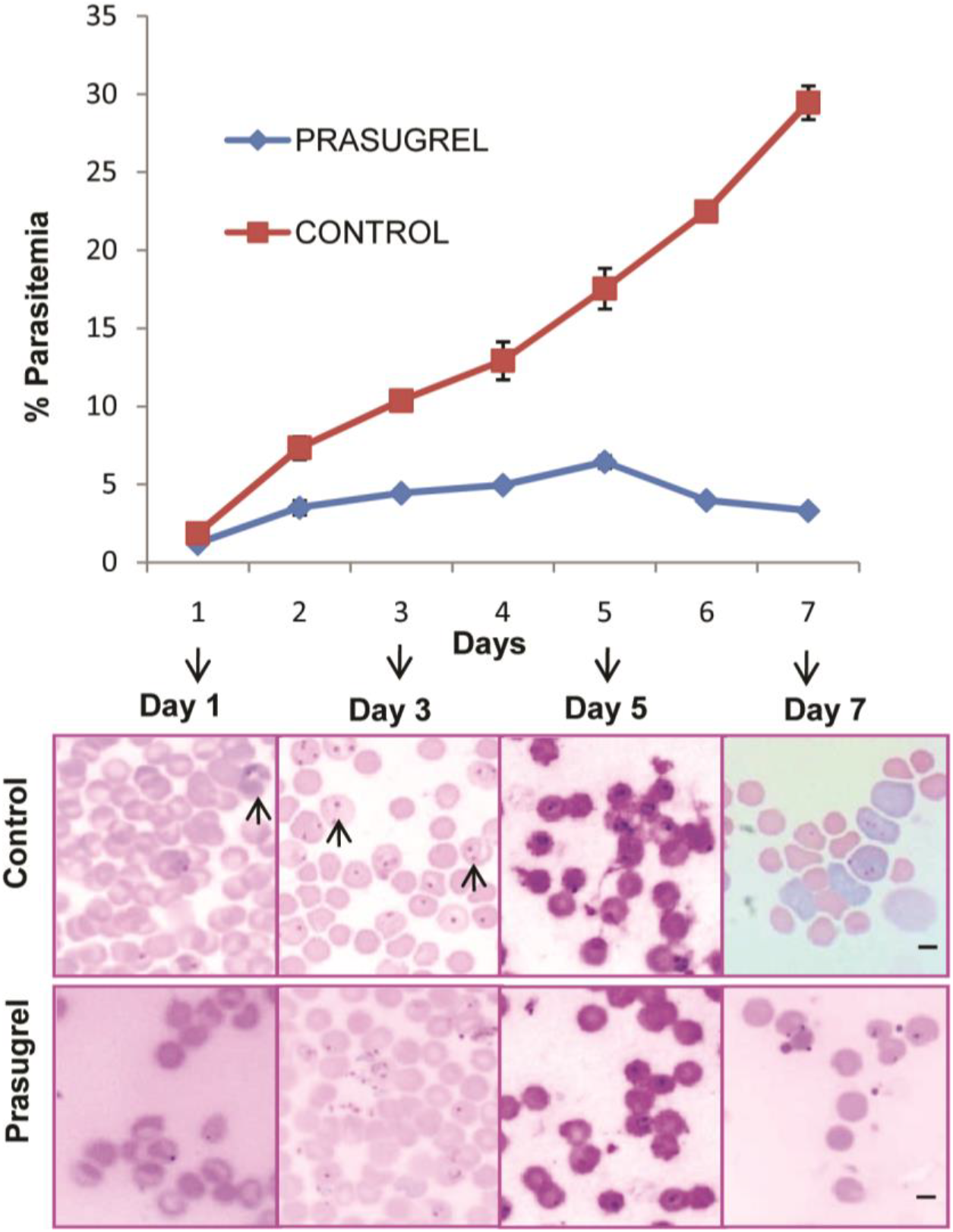
*In vivo* growth inhibitory effect of Prasugrel on *P. berghei* in mouse model. The percent parasitemia showed significant reduction in mice injected with Prasugrel as compared to vehicle control. The representative micrographs of Giemsa stained smears for both control and Prasugrel treated group are shown (bottom panel). Black arrows show parasitized RBCs. Scale= 5 μm.

### PfSRs are expressed in blood stages and localized to merozoites membrane

In order to delineate the functional role of PfSRs proteins, we have expressed the extracellular N-terminal regions of PfSRs as recombinant proteins in *E. coli* (Figure 4B(i)). Purified recombinant proteins rPfSR1_47_, rPfSR10_44_, rSR12_40_ and rSR25_35_ migrates on SDS-PAGE with an expected molecular weight of ~ 47 kDa, 44 kDa, 40 kDa and 35 kDa respectively (Figure 4B(ii)). Antibodies were raised against the individual recombinant proteins formulated with complete and incomplete Freund’s adjuvant in mice by standard immunization protocols. Immunofluorescence assays were carried out using the specific antisera to investigate stage specific expression of PfSRs in host erythrocytes. Analysis of confocal images taken at different time points showed PfSRs staining in late trophozoites, schizonts and merozoites but not in rings suggesting that the expression of these proteins initiates in the late trophozoite stage and then increases in mature schizonts and free merozoites (Figure 4A).

**Figure 4.**
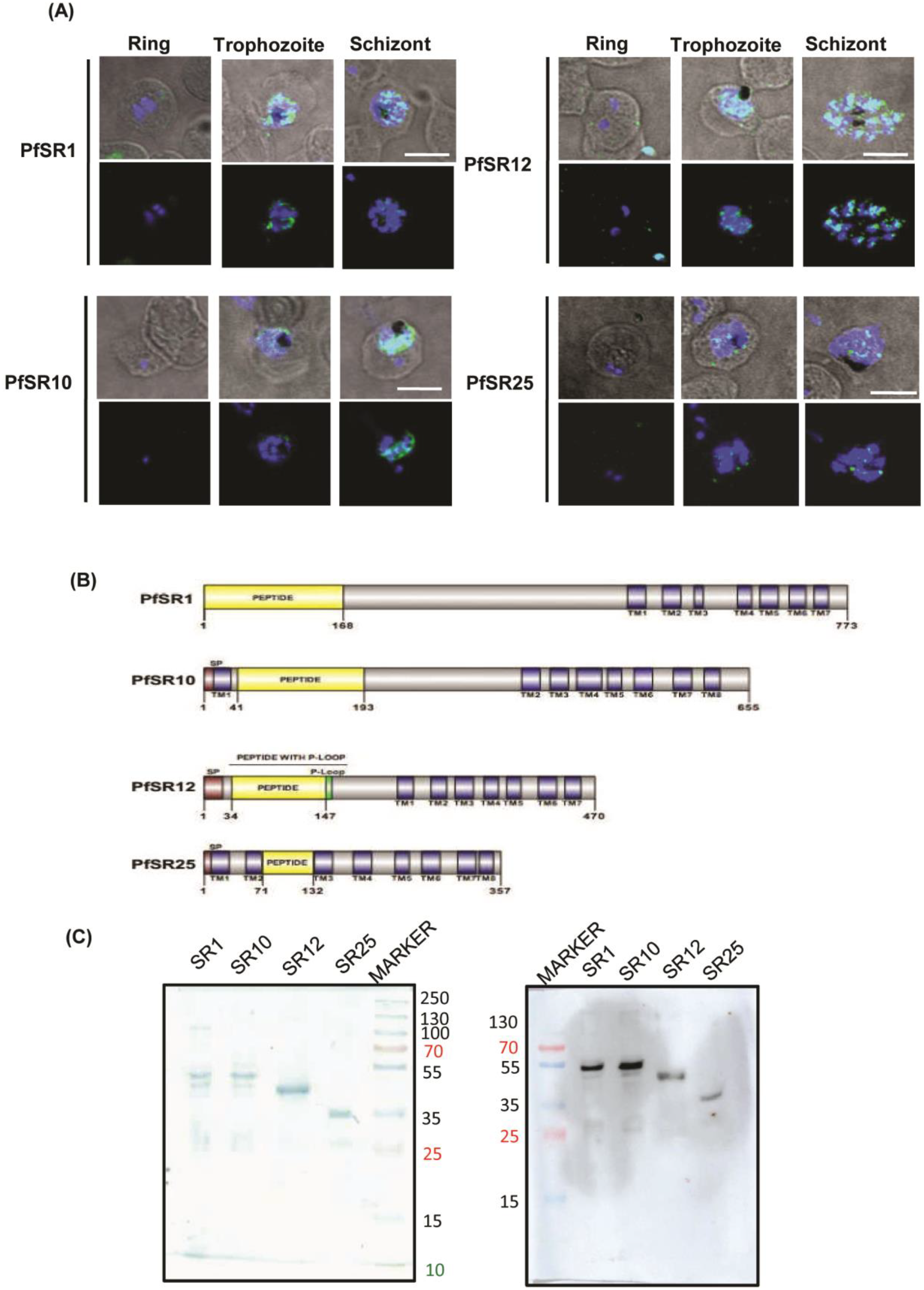
Protein structure and expression profile of *P. falciparum* serpentine receptors (PfSRs). (A) Stage-specific expression of PfSRs in the blood stages of *P. falciparum.* Expression of PfSRs starts in late trophozoite stage, with maximal expression in schizonts. Scale bar, 5 μm (B) (i) Schematic representation of four serpentine proteins showing functional domains (yellow) expressed as recombinant proteins. (ii) Purified recombinant proteins rPfSR1_47_, rPfSR10_44_, rSR12_40_ and rSR25_35_ with an expected molecular weight of ~ 47 kDa, 44 kDa, 40 kDa and 35 kDa respectively were separated by SDS-PAGE and detected by Coomassie staining. Western blotting with anti GST antibody confirms the purity of recombinant proteins.

For detailed studies on localization of PfSRs in schizonts and free merozoites, co-immunostaining was performed with known membrane marker protein i.e. merozoite surface protein 1 (MSP 1). Co-localization analysis with *Plasmodium* merozoite surface protein-1 (MSP-1) revealed that PfSRs are localized at parasite membrane (Figure 5). This suggests PfSRs might have an important role in membrane receptor signaling during invasion and egress.

**Figure 5.**
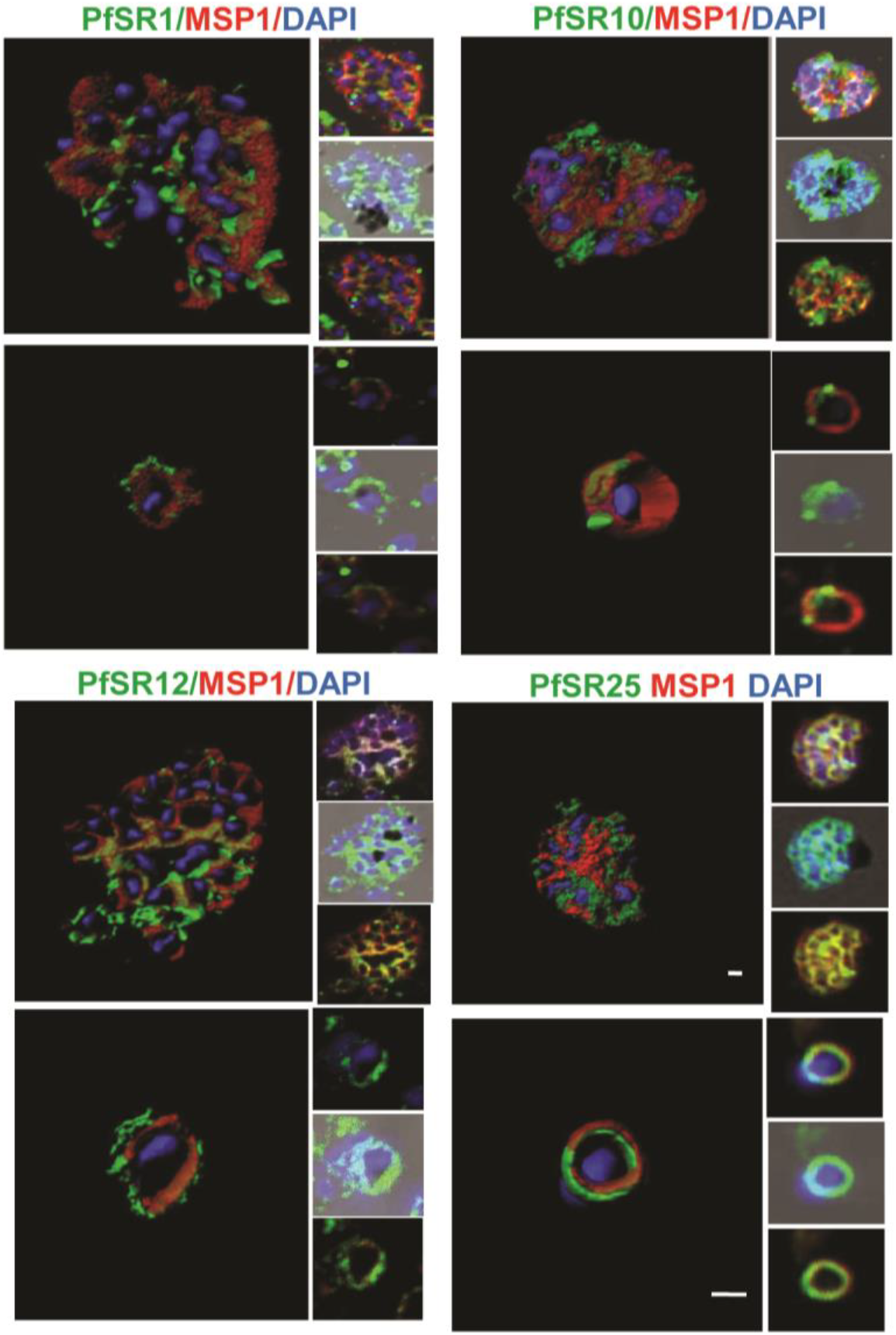
Localization of PfSRs in schizonts and merozoites of *P. falciparum.* Co-immunostaining of PfSRs (green) with merozoite surface protein MSP1 (red) revealed membrane localization of PfSRs in merozoites. Nuclei were counterstained with DAPI. Scale bar, 0.5 μm.

### PfSR12 is putative purinergic receptor which binds to ATP

To determine whether the predicted purinergic receptor PfSR12 is involved in purinergic signaling in parasite and indeed exhibits ATP binding properties, we performed ATP affinity assay using 6AH-ATP-Agarose beads with recombinant PfSR12 protein (rSR12_40_) and evaluated its specific binding with ATP. Elutes were run on SDS PAGE and protein band of rSR12_40_ corresponding to ~ 40 kDa was detected by silver staining (Figure 6A). No protein band corresponding to recombinant protein was observed in control beads lacking ATP. The results demonstrate that PfSR12 binds to ATP. Binding specificity was further evaluated by competition assay with free ATP or ADP. Western blot with anti rSR12_40_ sera showed that as compared with ADP, free ATP binds more effectively to the recombinant protein with 6AH-ATP beads in dose dependent manner suggesting specific binding of ATP with rSR12_40_ (Figure 6B). Prasugrel, a potent antagonist exerts its function by irreversibly binding to P2Y type purinergic receptor ^23^. To determine that PfSR12 is putative purinergic receptor expressed in blood stages of Plasmodium, we checked interaction of Prasugrel with PfSR12, by dose dependent titration of Prasugrel with rSR12_40_ in presence of ATP. We found that binding of recombinant PfSR12 with ATP beads was reduced at higher concentration of Prasugrel (Figure 6B). Together our results indicate PfSR12 as putative purinergic receptor and can be targeted by purinoreceptor antagonist Prasugrel.

**Figure 6.**
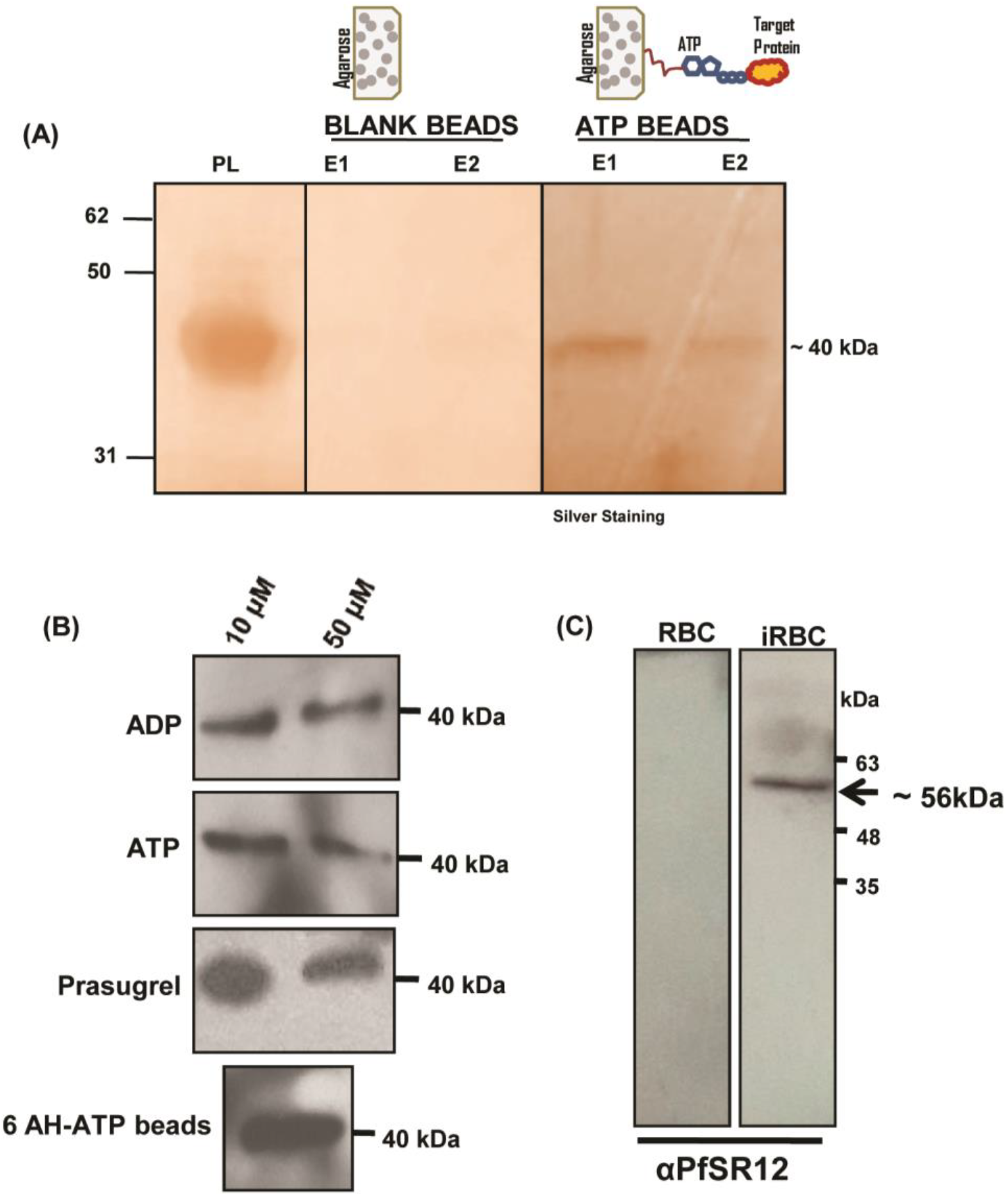
PfSR12 is purinergic receptor and expressed in late blood stages of *P. falciparum*. Binding of rSR12_40_ with ATP was analyzed by affinity assays using ATP beads. Eluted fraction (EL) showed a specific band of rSR12_40_ about ~ 40 kDa as detected by (A) silver staining and (B) Western blotting with polyclonal anti-rSR12_40_ mouse serum. No band was detected by preimmune serum (data not shown). No band was detected in blank agarose beads lacking ATP. PL: Preload (initial input of recombinant protein). (C) Blood stages such as late schizonts and merozoites were saponin treated, lysed using radioimmunoprecipitation assay (RIPA) buffer containing 1 % IGEPAL and 0.5 % sodium deoxycholate and run on SDS-PAGE. The presence of PfSR12 in lysate was detected by Western blotting using anti mouse serum. The band of expected size ~ 56 kDa of PfSR12 was detected in the parasite lysate. Uninfected RBC lysate was also probed with antisera of rSR12_40_ as a negative control.

### Detection of native PfSR12 by western blot

Furthermore, we confirmed the expression of native full length PfSR12 in *P. falciparum* 3D7 culture lysate by probing with anti-rSR12_40_ mouse sera (Figure 6C). A band of ~ 56 kDa, corresponding to full length size PfSR12 was detected in parasite lysate by Western blotting, thereby confirming the protein expression of putative purinergic receptor PfSR12 in the intraerythrocytic stages.

## Discussion

Signal transduction is a major mechanism regulating survival of malaria parasite and its development inside host erythrocytes ^7,24^. Studies have been carried out to elucidate the role of purinergic signaling in erythrocyte invasion by malaria parasite ^10,14^. There are conclusive evidences in the favour of existence of purinergic receptors in Plasmodium linked to increase in calcium, essential for invasion by malaria parasite ^6,25^. However in depth investigations for the expression of parasite receptors and associated signalling mechanisms for these phenotypes need to be explored further.

The present study shows Prasugrel, a potent thienopyridine class FDA approved prodrug and human P2Y_12_ receptor antagonist that inhibits the growth and development of Plasmodium in red blood cells. Growth inhibition assays showed prodrug Prasugrel has potent antimalarial efficacy than its active metabolite R-138727 or Suramin, a known non selective purinergic receptor antagonist. Intraerythrocytic life cycle of *P. falciparum* is quite synchronized *in vivo* ^26,27^ and it is important to understand at what stage of parasite’s life cycle, purinergic antagonist has maximal effect. Progression studies indicated the inhibitory effect of Prasugrel in late erythrocyte stages predominantly in the schizonts. Cruz *et al.* 2012 showed that purinergic antagonist modulates mouse malaria parasite *P. berghei* intraerythrocytic development and inhibit invasion. We also looked for intracellular growth inhibition of *P. berghei* in mouse model using Prasugrel. As it is already known that prodrug Prasugrel gets transformed into pharmacologically active metabolite, R-138727 *in vivo*. Therefore we presumed that inhibitory effect of Prasugrel against *P. berghei* in our *in vivo* studies is due to its active metabolite, however detailed studies need to be performed to validate this assumption. Our *in vivo* results confirmed growth inhibitory potential of Prasugrel as shown by *in vitro* studies.

In the attempt to identify purinergic receptors in malaria parasite that might regulate invasion and growth of parasite in host erythrocytes, we analyzed PfSRs, a class of GPCR like proteins present in *P. falciparum*, *P. berghei* and *P. yoelii* ^9^. In our previous work, we have shown ATP binding properties of PfSR12 and suggested as putative purinergic receptor by *in silico* studies ^12^. Based on our initial prediction, we expressed a 40 kDa N-terminal extracellular region of PfSR12 (rPfSR12_40_) with P-loop motif as recombinant protein and confirmed that it indeed binds to nucleotides like ATP with high specificity based on pull down assays with ATP beads. Interestingly the binding of recombinant protein with ATP beads was reduced upon addition of free ATP or Purinergic antagonist Prasugrel indicating specificity of recombinant protein for these molecules. However these results should be further confirmed experimentally by biochemical and biophysical approaches.

In this study, we also demonstrate by western blotting and IFA using specific antisera that PfSR12 is expressed in the blood stages of *P. falciparum* and its expression is localized to parasite membrane.

Preliminary docking studies indicated that all purinergic inhibitors used in our studies including Prasugrel, Suramin and R-138727 bind to PfSR12 protein in a thermodynamically favourable manner. The binding is majorly driven by Hydrogen bonds however Prasugrel association with PfSR12 is additionally stabilized by hydrophobic interactions, which act as predominant factor in protein stability. The stable complex formation between PfSR12 and purinergic receptor antagonists suggests its function as purinergic receptor in *P. falciparum*.

Taken together, this study reiterates the presence of purinergic receptor in blood stages of malaria parasite and the effect of P2Y purinergic receptor inhibitors on growth and development of asexual stages in erythrocytes.

Our findings advocate purinergic signaling as a key target for designing novel antimalarial agents with the potential to block growth and invasion of parasite in host erythrocytes. The known purinoreceptor blocker, Suramin has been shown to inhibit *P. falciparum* invasion into host erythrocytes and is commonly used to treat parasitic diseases in humans including African sleeping sickness ^28,29^. However this drug is not suitable for any chemotherapeutic purposes due to toxicity and severe side effects. Our work justifies further analysis of Prasugrel (platelet aggregation inhibitor drug) as a valid option of repositioning in malaria, a dreadful infectious disease. Drug Repositioning has been employed to significantly shorten the drug discovery time for many diseases including malaria ^30^. Furthermore our work identifies PfSR12 could be putative purinergic receptor in parasite and novel drug target for development of antimalarials.

## Acknowledgements

We are grateful to Advanced Instrumentation Research Facility (AIRF), Jawaharlal Nehru University for confocal microscopy, Central Instrumentation Facility (CIF) of SCMM, JNU for flow cytometry and Animal House facility for animal experiments. The lab facility of Shiv Nadar University is also acknowledged.

## Author’s contribution

SG and SS conceived and designed the experiments. SG, NJ and MS performed laboratory experiments. SG analyzed the data and wrote the manuscript. NJ did bioinformatics work. SG and SS drafted and critically revised the manuscript.

## References

1 Iyer, J. K., Amaladoss, A., Genesan, S. & Preiser, P. R. Variable expression of the 235 kDa rhoptry protein of Plasmodium yoelii mediate host cell adaptation and immune evasion. Molecular microbiology 65, 333–346, doi:10.1111/j.1365-2958.2007.05786.x (2007).

2 Singh, S., Alam, M. M., Pal-Bhowmick, I., Brzostowski, J. A. & Chitnis, C. E. Distinct external signals trigger sequential release of apical organelles during erythrocyte invasion by malaria parasites. PLoS pathogens 6, e1000746, doi:10.1371/journal.ppat.1000746 (2010).

3 Schwiebert, E. M. & Zsembery, A. Extracellular ATP as a signaling molecule for epithelial cells. Biochimica et biophysica acta 1615, 7–32 (2003).

4 Burnstock, G. Introduction: P2 receptors. Current topics in medicinal chemistry 4, 793–803 (2004).

5 Ellsworth, M. L. Red blood cell-derived ATP as a regulator of skeletal muscle perfusion. Medicine and science in sports and exercise 36, 35–41, doi:10.1249/01.MSS.0000106284.80300.B2 (2004).

6 Levano-Garcia, J., Dluzewski, A. R., Markus, R. P. & Garcia, C. R. Purinergic signalling is involved in the malaria parasite Plasmodium falciparum invasion to red blood cells. Purinergic signalling 6, 365–372, doi:10.1007/s11302-010-9202-y (2010).

7 Cruz, L. N. et al. Extracellular ATP triggers proteolysis and cytosolic Ca(2)(+) rise in Plasmodium berghei and Plasmodium yoelii malaria parasites. Malaria journal 11, 69, doi:10.1186/1475-2875-11-69 (2012).

8 Fleck, S. L. et al. Suramin and suramin analogues inhibit merozoite surface protein-1 secondary processing and erythrocyte invasion by the malaria parasite Plasmodium falciparum. The Journal of biological chemistry 278, 47670–47677, doi:10.1074/jbc.M306603200 (2003).

9 Madeira, L. et al. Genome-wide detection of serpentine receptor-like proteins in malaria parasites. PloS one 3, e1889, doi:10.1371/journal.pone.0001889 (2008).

10 Lindner, S. E. et al. Total and putative surface proteomics of malaria parasite salivary gland sporozoites. Molecular & cellular proteomics: MCP 12, 1127–1143, doi:10.1074/mcp.M112.024505 (2013).

11 Moraes, M. S. et al. Plasmodium falciparum GPCR-like receptor SR25 mediates extracellular K(+) sensing coupled to Ca(2+) signaling and stress survival. Scientific reports 7, 9545, doi:10.1038/s41598-017-09959-8 (2017).

12 Subudhi, A. K. et al. Malaria parasites regulate intra-erythrocytic development duration via serpentine receptor 10 to coordinate with host rhythms. Nature communications 11, 2763, doi:10.1038/s41467-020-16593-y (2020).

13 Gupta, S., Singh, D. & Singh, S. In silico characterization of Plasmodium falciparum purinergic receptor: a novel chemotherapeutic target. Systems and synthetic biology 9, 11–16, doi:10.1007/s11693-015-9165-y (2015).

14 Jensen, J. B. In vitro culture of Plasmodium parasites. Methods in molecular medicine 72, 477–488, doi:10.1385/1-59259-271-6:477 (2002).

15 Dery, V. et al. An improved SYBR Green-1-based fluorescence method for the routine monitoring of Plasmodium falciparum resistance to anti-malarial drugs. Malaria journal 14, 481, doi:10.1186/s12936-015-1011-x (2015).

16 Xu, D. & Zhang, Y. Improving the physical realism and structural accuracy of protein models by a two-step atomic-level energy minimization. Biophysical journal 101, 2525–2534, doi:10.1016/j.bpj.2011.10.024 (2011).

17 Spessard, G. O. ACD Labs/LogP dB 3.5 and ChemSketch 3.5. J. Chem. Inf. Comput. Sci., doi:10.1021/ci980264t (1998).

18 Cousins, K. R. Computer review of ChemDraw Ultra 12.0. Journal of the American Chemical Society 133, 8388, doi:10.1021/ja204075s (2011).

19 Cousins, K. R. Computer Review of ChemDraw Ultra 12.0 ChemDraw Ultra 12.0. CambridgeSoft, 100 Cambridge Park Drive, Cambridge, MA 02140. J. Am. Chem. Soc., doi:10.1021/ja204075s (2011).

20 Salentin, S., Schreiber, S., Haupt, V. J., Adasme, M. F. & Schroeder, M. PLIP: fully automated protein-ligand interaction profiler. Nucleic acids research 43, W443–447, doi:10.1093/nar/gkv315 (2015).

21 Racine, J. The Cygwin tools: a GNU toolkit for Windows. J. Appl. Econom., doi:10.1002/1099-1255(200005/06)15:3<331::aid-jae558>3.0.co;2-g (2000).

22 Wickremsinhe, E. R. et al. Stereoselective metabolism of prasugrel in humans using a novel chiral liquid chromatography-tandem mass spectrometry method. Drug metabolism and disposition: the biological fate of chemicals 35, 917–921, doi:10.1124/dmd.106.014530 (2007).

23 Algaier, I., Jakubowski, J. A., Asai, F. & von Kugelgen, I. Interaction of the active metabolite of prasugrel, R-138727, with cysteine 97 and cysteine 175 of the human P2Y12 receptor. Journal of thrombosis and haemostasis: JTH 6, 1908–1914, doi:10.1111/j.1538-7836.2008.03136.x (2008).

24 Hotta, C. T. et al. Calcium-dependent modulation by melatonin of the circadian rhythm in malarial parasites. Nature cell biology 2, 466–468, doi:10.1038/35017112 (2000).

25 Tanneur, V. et al. Purinoceptors are involved in the induction of an osmolyte permeability in malaria-infected and oxidized human erythrocytes. FASEB journal: official publication of the Federation of American Societies for Experimental Biology 20, 133–135, doi:10.1096/fj.04-3371fje (2006).

26 Fidock, D. A., Rosenthal, P. J., Croft, S. L., Brun, R. & Nwaka, S. Antimalarial drug discovery: efficacy models for compound screening. Nature reviews. Drug discovery 3, 509–520, doi:10.1038/nrd1416 (2004).

27 Kesely, K. R., Pantaleo, A., Turrini, F. M., Olupot-Olupot, P. & Low, P. S. Inhibition of an Erythrocyte Tyrosine Kinase with Imatinib Prevents Plasmodium falciparum Egress and Terminates Parasitemia. PloS one11, e0164895, doi:10.1371/journal.pone.0164895 (2016).

28 Ullmann, H. et al. Synthesis and structure-activity relationships of suramin-derived P2Y11 receptor antagonists with nanomolar potency. Journal of medicinal chemistry 48, 7040–7048, doi:10.1021/jm050301p (2005).

29 Abbracchio, M. P. et al. International Union of Pharmacology LVIII: update on the P2Y G protein-coupled nucleotide receptors: from molecular mechanisms and pathophysiology to therapy. Pharmacological reviews 58, 281–341, doi:10.1124/pr.58.3.3 (2006).

30 Nzila, A., Ma, Z. & Chibale, K. Drug repositioning in the treatment of malaria and TB. Future medicinal chemistry 3, 1413–1426, doi:10.4155/fmc.11.95 (2011).

